# Contact lenses can cause the reverse Pulfrich effect and anti-Pulfrich monovision corrections can eliminate it

**DOI:** 10.1101/2020.04.05.026534

**Authors:** Victor Rodriguez-Lopez, Carlos Dorronsoro, Johannes Burge

## Abstract

Interocular differences in image blur can cause dramatic misperceptions of the distance and three-dimensional direction of moving objects. This new illusion—the reverse Pulfrich effect—is caused by the optical conditions induced by monovision, a common correction for presbyopia. Fortunately, anti-Pulfrich monovision corrections, in which the blurring lens is slightly darkened, can eliminate the illusion for a wide range of viewing conditions. However, the reverse Pulfrich effect and the efficacy of anti-Pulfrich corrections have previously been demonstrated only with trial lenses. This situation should be addressed, for both clinical and scientific reasons. First, monovision is most commonly prescribed with contact lenses. It is important to replicate these effects in the most common monovision delivery system. Second, trial lenses of different powers, unlike contacts, cause large magnification differences between the eyes. To confidently attribute the reverse Pulfrich effect to differences in optical blur between the eyes, and to ensure that the reported effect sizes are reliable, one must control for magnification. Here, in a within observer study with five separate experiments, we demonstrate i) that contact lenses induce reverse Pulfrich effects that are indistinguishable from those induced by trial lenses, ii) that overall magnification differences do not cause or impact the Pulfrich effect, and iii) that anti-Pulfrich corrections (i.e. darkening the blurring lens) are equally effective when induced by contact lenses and by trial lenses.

## Introduction

Monovision is a widely prescribed correction for presbyopia, the age-related loss of focusing ability, impacting 2 billion people worldwide (Charman, 2008; Fricke et al., 2018). Monovision corrections serve as an alternative to reading glasses, bifocals, and progressive lenses for millions of people (Bennett, 2008; Morgan, Efron, & Woods, 2011). With monovision, each eye is fit with a lens that sharply focuses light from a different distance. One eye is corrected for ‘far vision’, while the other eye is corrected for ‘near vision’. Monovision thus intentionally induces differences in image blur between the eyes. The eye corrected for near vision will form sharper images of near objects like books or mobile phones, and blurrier images of far objects like mountains or street signs. Similarly, the eye corrected for far vision will form blurrier images of near objects and sharper images of far objects.

Burge et al. (2019) recently reported that interocular differences in optical blur, like those induced by monovision corrections, have the potential to cause large errors in estimating the distance and 3D direction of moving objects. Under some conditions, the perceptual errors may be large enough to impact public safety. For example, the distance to a cyclist in cross-traffic may be overestimated by nearly 9ft, the width of a narrow lane of traffic (Fig. 1A; Burge, Rodriguez-Lopez, & Dorronsoro, 2019). The illusion occurs because the image in the blurrier eye is processed more quickly by a few milliseconds than the image in the sharper eye. For moving targets, this interocular mismatch in processing speed causes a neural disparity, which results in the misperceptions (see Results). The new illusion—the reverse Pulfrich effect—is closely related to the classic Pulfrich effect, a well-known stereo-motion phenomenon that was first discovered 100 years ago.

**Figure 1.**
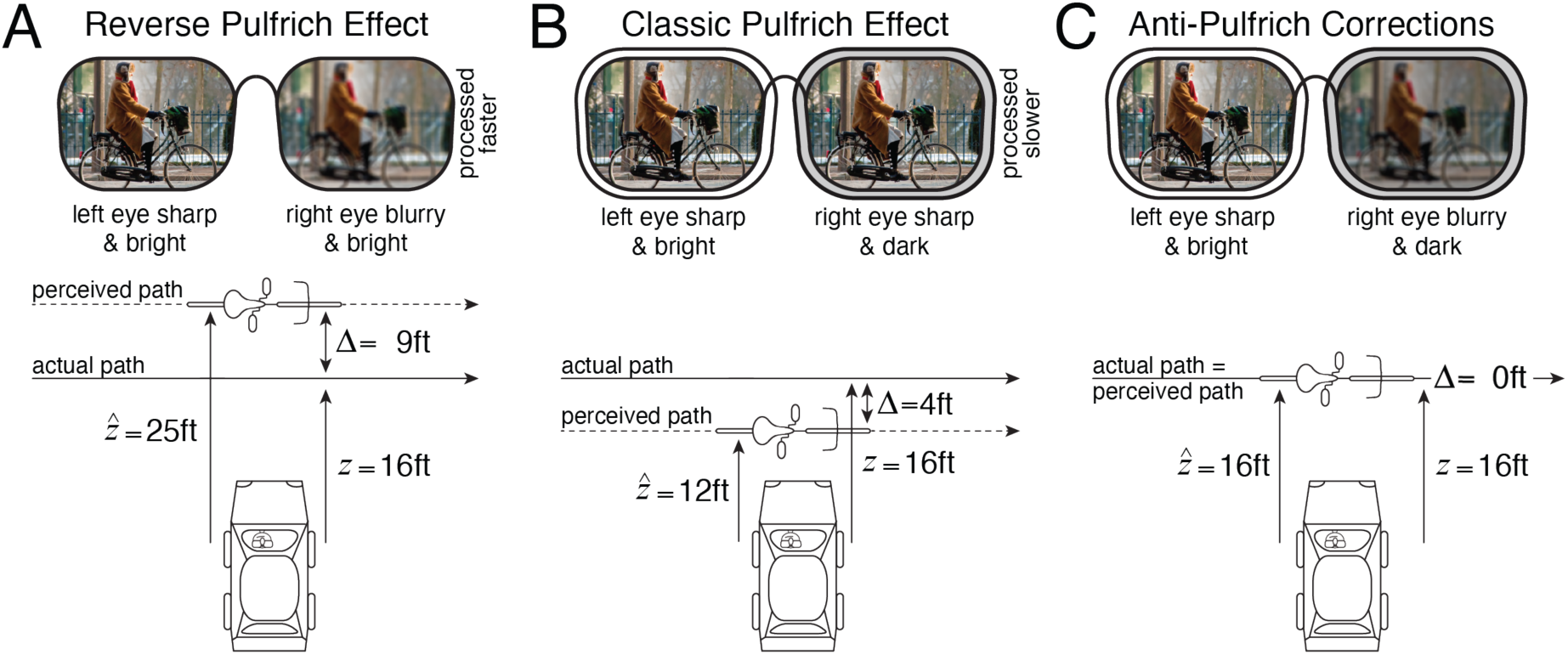
Reverse Pulfrich effect, classic Pulfrich effect, and anti-Pulfrich monovision corrections. **A** Interocular blur differences like those induced by monovision corrections cause the ‘reverse Pulfrich effect’, a substantial misperception of the distance of moving objects. If the left eye is sharp and the right eye is blurred, an object moving from left to right will be misperceived as farther away than it actually is, and vice versa. The blurry image is processed faster than the sharp image, causing a neural disparity which leads to the depth misperceptions. In some scenarios, the distance misestimates can be substantial. Burge et al. (2019) reported that, for an individual observer with a typical monovision correction strength of 1.5D, a cyclist moving left to right at 15mph at a distance of 16ft may be estimated to be at 25ft. This overestimation of 9ft is approximately the width of a narrow lane of traffic. These misperceptions occur because the blurrier eye is processed more quickly by only a few milliseconds (see Results). **B** Interocular luminance differences cause the classic Pulfrich effect. When both eyes are sharp, the darker eye is processed slower. If the left eye is bright and the right eye is dark, and both eyes are sharply focused, the distance to the same cyclist will be underestimated, because the darker eye is processed more slowly by a few milliseconds. **C** Anti-Pulfrich monovision corrections can eliminate the misperceptions by darkening the blurring lens. The reverse and classic Pulfrich effects cancel each other out.

The classic Pulfrich effect is induced by an interocular difference in retinal illuminance. However, darkening the image in one eye has the opposite effect of blur (Pulfrich, 1922). The darker image is processed more slowly rather than more quickly (Lit, 1949; Rogers & Anstis, 1972; Wilson & Anstis, 1969). The resulting illusions are thus similar to the illusions associated with the reverse Pulfrich effect, except that the classic Pulfrich effect causes distance underestimation where the reverse Pulfrich effect causes distance overestimation, and vice versa (Fig. 1B).

Burge et al. (2019) also demonstrated that anti-Pulfrich monovision corrections can eliminate the depth and motion misperceptions associated with the reverse and classic Pulfrich effects. The logic behind anti-Pulfrich corrections is simple. Blurry images are processed more quickly than sharp images (i.e. the reverse Pulfrich effect). Dark images are processed more slowly than bright images (i.e. the classic Pulfrich effect). Thus, if the blurry image is darkened appropriately, the two differences in processing speed should cancel one another out and eliminate the misperceptions (Fig. 1C). To date, however, the efficacy of anti-Pulfrich monovision corrections has been demonstrated only with interocular differences in optical power induced by trial lenses (Burge et al., 2019). Monovision prescriptions are most commonly prescribed with contact lenses (Evans, 2007). (Surgically implanted interocular lenses are second most common (Davidson et al., 2016; Wolffsohn & Davies, 2019; Xiao, Jiang, & Zhang, 2011).) Hence, it is important to demonstrate that anti-Pulfrich corrections are effective when implemented with the ophthalmic corrections that are most relevant to clinical practice.

Another reason to study the reverse Pulfrich effect and anti-Pulfrich corrections with contact lenses is that, unlike trial lenses, contacts do not introduce retinal magnification differences between the eyes. The retinal magnification induced by a lens increases with its optical power and its distance from the eye (see Methods). Trial lenses, like eyeglasses, are positioned a considerable distance from the eye. As a consequence, trial lenses of different powers induce both blur and magnification differences. The original demonstration of the reverse Pulfrich effect thus leaves open the possibility that the reverse Pulfrich could be impacted by interocular differences in magnification, rather than to interocular differences in blur. Although theoretical considerations and previously performed control experiments make this possibility unlikely, it should still be established empirically that differences in image magnification play no role in driving the effect.

A typical monovision correction induces a 1.5D difference in optical power between the eyes (Evans, 2007). For contacts, which are fit directly on the cornea, this power difference translates into a magnification difference between the eyes of only 0.4% (Fig. 2A; see Methods). For eyeglasses, which are typically positioned 10-14mm from the cornea, the same power difference translates into a magnification difference of between 1.5% and 2.1% (Fig. 2B). Magnification differences of this size are thought to cause visual discomfort and other clinical issues (Bannon, Neumueller, Boeder, & Burian, 1970). This is the reason that monovision is most often implemented with contact lenses and with surgically implanted interocular lenses, both of which induce negligible magnification differences.

**Figure 2.**
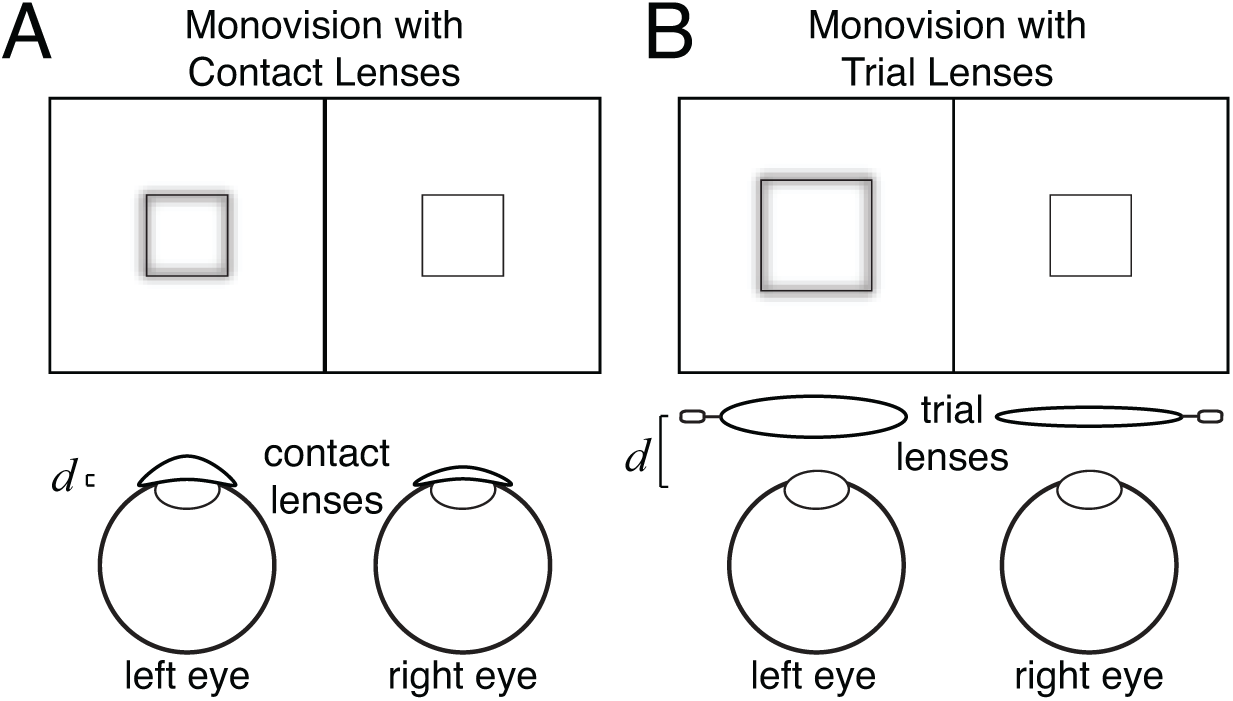
Magnification differences with contacts and trial lenses. **A** Contact lenses with different power in the two eyes create interocular differences in blur with negligible interocular differences in magnification. Contact lenses produce negligible image magnification because they are fit directly on the cornea. The distance *d* between the contact lens and the entrance pupil of the eye is quite small. **B** Trial lenses with different powers in the two eyes create interocular differences in blur with non-negligible magnification differences between the eyes. Trial lenses produce substantially more image magnification than contact lenses of the same power because the distance between the lens and the entrance pupil is considerably larger (i.e. 10-14mm).

Measuring the reverse Pulfrich effect with soft contact lenses has the benefit of i) isolating the impact of blur from the potential impact of magnification on processing speed, ii) testing for misperceptions in the optical conditions most similar to those induced by eye care practitioners, and iii) advancing towards a clinically applicable anti-Pulfrich correction.

## Methods

### Participants

Two male and two female observers between the ages of 25 and 30 participated in the experiment. One male observer was an author; all other observers were naïve to the purposes of the experiment. Anisometropia, a difference in refractive error between the eyes, was equal to or lower than 0.5D in all observers. The amount of astigmatism was subclinical in all observers but one, who had astigmatism of 0.5D. Visual acuity was normal or corrected-to-normal in both eyes of each observer. Stereo-acuity was also normal, as assessed with the Titmus stereo test.

### Apparatus

The stimulus was displayed on a stereo-3D UK UHD 49” monitor (LG49UH850V, LG). The monitor uses vertical spatial interlacing (e.g. pixels in even rows to the left eye & pixels in odd rows to the right eye) to present different temporally coincident images to the left and right eyes. The maximum luminance of the monitor was 400cd/m^2^. The monitor was driven by a NVIDIA® Quadro® P4000 dual Graphic card. The refresh rate was 60Hz (i.e. 60Hz/eye).

Passive circular polarization glasses selectively passed the appropriate image to each eye. The spatial resolution of the display was 3840×2160 pixels. After filtering by the glasses, only 3840×1080 interlaced pixels reached each eye. Combined with a transmittance of slightly less than 1.0, the effective luminance of the monitor for each eye was slightly less than 200cd/m^2^.

The observer viewed the monitor from a distance of 2m, with his/her head stabilized by a chin and forehead rest. At this viewing distance, each pixel subtended 0.46 arcmin of visual angle. Observers viewed the display through custom-built mounts for trial lenses.

The mounts were horizontally and vertically adjusted so that the optical element was centered along the line of sight of each eye.

### Stimuli

The target stimulus was a 0.25×1.00° white vertical bar that oscillated horizontally on a gray background (Fig. 3A). The target bar traversed one cycle of a cosinusoidal trajectory during each trial. The left- and right-eye onscreen bar positions in degrees of visual angle were given by

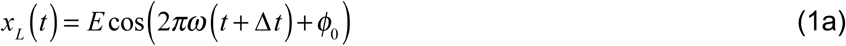

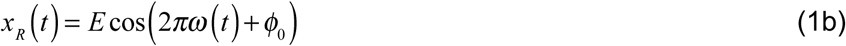

where *E* is the target movement amplitude in degrees of visual angle, *ω* is the temporal frequency of the target movement, *ϕ*_0_ is the starting phase, *t* is time in seconds, and Δ*t* is the onscreen delay between the left- and right-eye target images. The onscreen interocular delay between the left- and right-eye target images controlled whether stereo information specified ‘front left’ or ‘front right’ motion (Fig. 3B). The onscreen binocular disparity associated with a given interocular delay as a function of time is given by

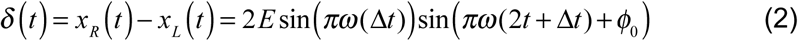

where negative disparities are crossed (i.e. nearer than the screen) and positive disparities are uncrossed (i.e. farther than the screen).

**Figure 3.**
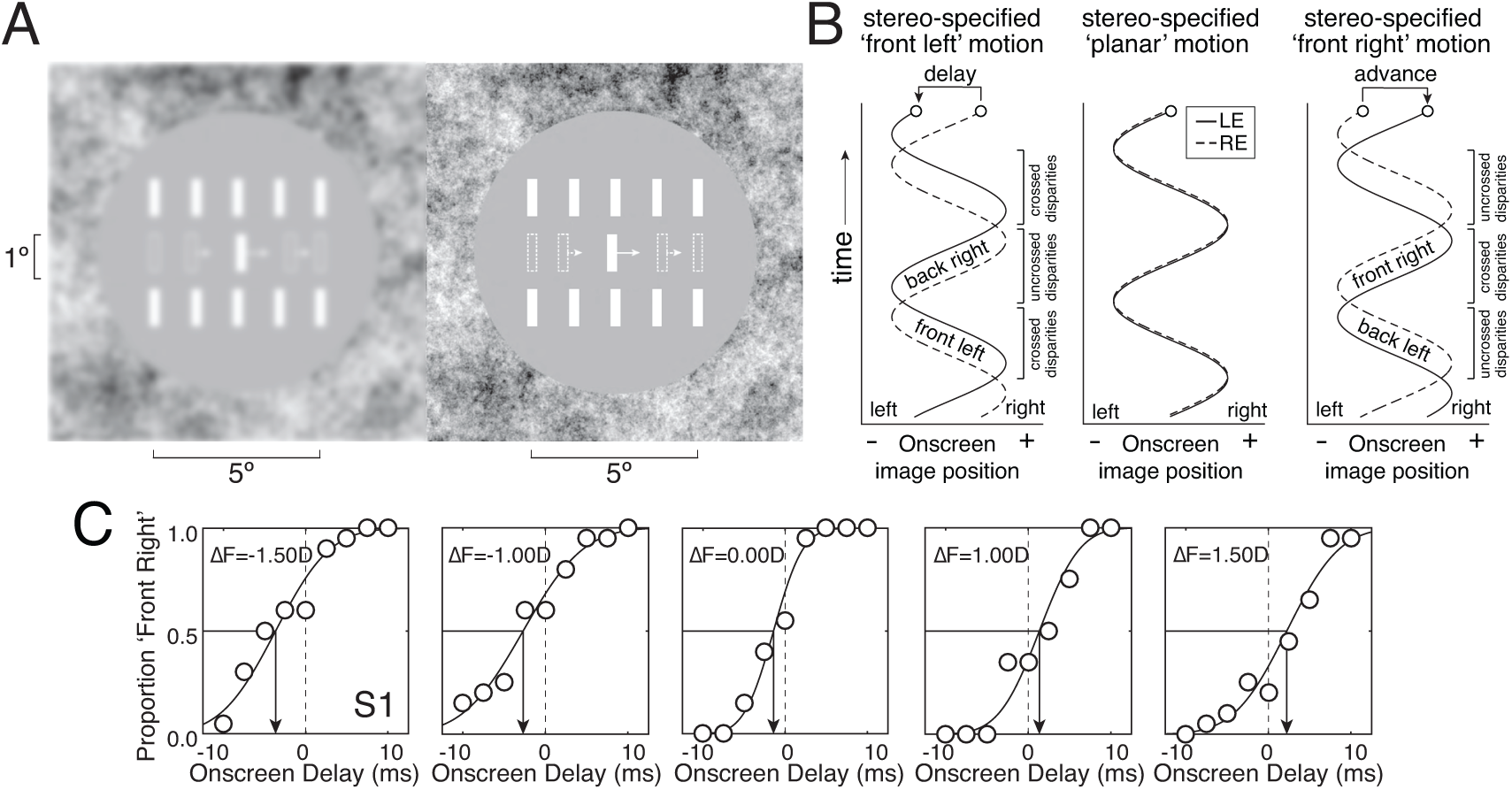
Binocular stimulus, time-course of stimulus presentation, and psychometric functions. **A** The target was a dichoptically presented horizontally moving white bar. The left-eye image is blurred to simulate the optical blur that was induced in the experiment; no onscreen blur was present in this experiment. White arrows show target motion, speed, and direction. Dashed bars show example stimulus positions throughout a trial. Arrows and dashed bars are both for illustrative purposes and were not present in the actual stimulus. Fuse the two half-images to perceive the target bar in 3D on one frame of the movie. Cross-fusers will see a depiction of ‘front right’ motion on this frame. Divergent-fusers will see ‘back right’ motion on this frame and would answer ‘front left’ for the complete one-cycle trial. **B** Left-eye and right-eye onscreen horizontal image positions as a function of time (solid and dashed curves, respectively) when the left-eye image was delayed onscreen, coincident with, or advanced onscreen relative to the right-eye image. **C** The task was to report whether the target bar appeared to be moving ‘front left’ or ‘front right’ with respect to the screen. Psychometric functions for the first human observer as a function of onscreen delay in five conditions in Exp. 1. Each condition had a different interocular difference in focus error (i.e. Δ*F*=[−1.5D, −1.0D, 0.0D, 1.0D, 1.5D]). The point of subjective equality (PSE, black arrows) changes systematically with the difference in focus error, indicating that the difference in focus error systematically impacts the neural differences in processing speed between the eyes.

The trial duration was one second. The movement amplitude was 2.5° of visual angle (i.e. 5.0° total change in visual angle in each direction during each trial). The starting position of the target bar was randomly selected on each trial to start 2.5° to the right (i.e. *ϕ*_0_ =0) or 2.5° to the left (i.e. *ϕ*_0_ = *π*) of the screen center. Throughout each trial, observers fixated a white fixation dot at the center of the screen. Two sets of five stationary white ‘picket fence’ bars that were identical to the target bar flanked the region of the screen traversed by the target bar. The picket fence bars served as a reference to the stereo-specified distance of the screen. The screen periphery was covered with a 1/f noise texture to aid binocular fusion; the 1/f noise texture also served as a reference to the stereo-specified screen distance. Stimuli were generated with Matlab (Mathworks, Inc.) and presented via PsychToolbox 3 (Brainard, 1997).

### Procedure

Each moving bar stimulus was presented as part of a one-interval two-alternative forced-choice (2AFC) procedure. The task was to report, via a keypress, whether the target bar was moving leftwards (‘front left’) or rightwards (‘front right’) when the bar appeared to be in front of the plane of the screen. Nine evenly spaced levels of onscreen interocular delay between −10 and 10 milliseconds were presented with the method of constant stimuli. Twenty trials per level were collected for a total of 180 trials per condition.

The proportion ‘front right’ responses were recorded as a function of onscreen interocular delay and fit with a cumulative Gaussian via maximum likelihood methods. The point of subjective equality (PSE) indicates the onscreen interocular delay necessary to make the target appear to move within the screen plane. The PSE is opposite in sign and equal in magnitude to the neural difference in processing speed between the eyes.

Data was collected from each human observer in five different experiments; each experiment had multiple conditions (see below). In a given experiment, data was collected across all conditions in counterbalanced blocks of 90 trials each. Each block took approximately 2.5 minutes to complete. Experiment 1 measured the impact of interocular blur differences induced by contact lenses. Experiment 2 measured the impact of interocular blur differences induced by trial lenses. Experiment 3 measured the impact of interocular luminance differences between the eyes. Experiment 4 measured the efficacy of anti-Pulfrich monovision corrections with contact lenses. And Experiment 5 measured whether interocular magnification differences impact processing speed.

### Optical Conditions in the Experiments

To determine the power of the lenses that would be used to induce the desired focus error and the resulting optical blur in each eye, we performed three separate steps. First, we chose an optical power equal to the desired focus error (i.e. excess power) in one eye; the excess power was always positive. Second, we added a distance compensation power to set the optical distance of the screen to optical infinity; given that the actual distance of the screen was 2m, the required distance compensation power was +0.5D. Third, using standard methods of subjective refraction, we measured and corrected each observer to ensure that uncorrected refractive errors did not compromise the desired optical conditions. Importantly, adds of excess optical power to a screen already at optical infinity position the optical distance of the screen beyond optical infinity. Our procedure, therefore, renders accommodation unable to compensate for the desired focus error caused by the excess optical power.

The total optical power of the associated lens is thus given by

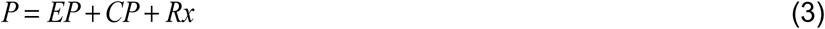

Where *EP* is the excess power (i.e. the desired focus error), *CP* is the compensation power, and *Rx* is the refractive error of the human observer, all of which are expressed in diopters. Combining all the total amount of optical power in one lens instead of using multiple lenses minimizes potential interocular differences due to reflections and transmission.

In experiments with contact lenses, the total optical power (Eq. 3) was delivered with ACUVUE Moist monofocal soft contact lenses (Johnson & Johnson Vision Care, Jacksonville, FL). In the experiment with trial lenses, the excess and compensation powers were delivered with a trial lens, and the refractive error of each human observer was compensated for by their own spectacles. In all other experiments, the total power was delivered by a single contact lens. The human observers had different amounts of myopia. Hence, across observers and experimental conditions (see below), the nominal power of the contact lenses ranged from −2.75D to +0.75D.

### Interocular differences in luminance

To induce the required differences in retinal illuminance, we reduced the luminance of the perturbed eye’s onscreen image by a scale factor equivalent to the transmittance of a neutral density filter with a particular ocular density. The transmittance is given by

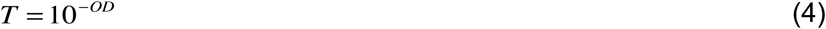

where *OD* is the optical density of the filter. We have previously verified that this procedure—using ‘virtual’ neutral density filters—yields results that are equivalent to using real neutral density filters (Burge et al., 2019).

### Image magnification

The relative magnification—the magnification caused by an ophthalmic lens relative to the naked eye—depends both on the power of the lens and on the distance of the lens to the entrance pupil of the eye (Bass, Enoch, & Lakshminarayanan, 2010). Under the thin lens approximation, which is appropriate in the present circumstances, image magnification is given by

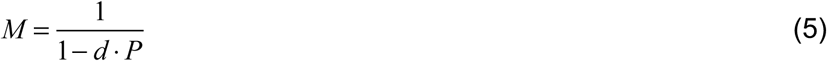

where *d* is the distance of the lens in meters to the entrance pupil and *P* is the power of the lens in diopters.

### Interocular differences in focus error, optical density, and magnification

The interocular difference in focus error (i.e. optical power) is defined as the difference in excess optical power between the eyes

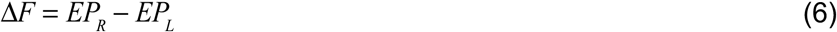

Where *EP*_*L*_ and *EP*_*R*_ are the excess optical powers in diopters of the left and right eyes, respectively. In experiments having conditions with non-zero differences in focus error (Exps. 1, 2, and 4), excess power (i.e. *EP* >0.0D) was induced in one eye only— the perturbed eye—while the other eye was kept sharply focused at the screen distance (i.e. *EP* =0.0D). In these experiments, the interocular difference in focus error ranged from −1.5D to 1.5D.

The interocular difference in luminance is quantified by the interocular difference in optical density

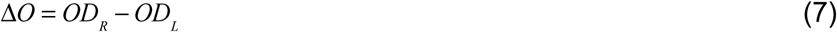

Where *OD*_*L*_ and *OD*_*R*_ are the optical density of the neutral density filters in the left and right eyes. In experiments having conditions with non-zero interocular differences in optical density (Exps. 3 and 4), the luminance in one eye was reduced whereas the other eye was left unperturbed. In these experiments, the interocular difference in optical density ranged from −0.15OD to 0.15OD. An optical density of 0.15OD corresponds to a transmittance of 70.8%.

The interocular difference in onscreen magnification is given by

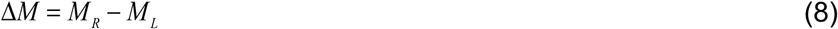

where *M*_*L*_ and *M*_*R*_ represent the magnification associated with the left- and right-eye images, respectively. In the experiment that manipulated magnification differences onscreen (Exp. 5), the interocular difference in magnification ranged from −3.6% to 3.6%. Magnification differences of this size are twice the magnification difference induced by trial lenses differing by 1.5D in optical power.

### Quantifying differences in processing speed from differences in blur and luminance

The interocular difference in processing speed was measured for interocular differences in focus error ranging from −1.5D to 1.5D and interocular differences in optical density ranging from −0.15OD to −0.15OD. The interocular difference in processing speed (i.e. interocular delay) is linearly related to the interocular difference in focus error

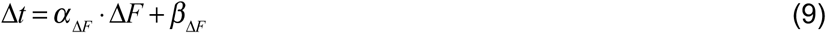

where *α*_Δ*F*_ and *β*_Δ*F*_ are the slope and intercept of the best line fit via least squared regression (Burge et al., 2019). Just as with blur differences, the interocular delay is linearly related to the difference between the optical densities of the virtual filters in the two eyes. Specifically,

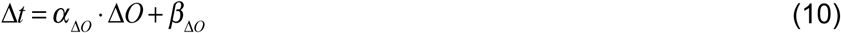

Δ*O* is the interocular difference in optical density, and *α* _Δ*O*_ and *β*_Δ*O*_ are the slope and the intercept of the best line fit via linear regression. Linear regression was similarly used to fit the pattern of interocular delays with interocular differences in magnification.

### Anti-Pulfrich monovision corrections

The interocular differences in processing speed caused by a unit difference in optical power (i.e. *α* _Δ*F*_) and a unit difference in optical density (i.e. *α* _Δ*O*_) can be used to determine the luminance difference required to null processing speed differences caused by an arbitrary difference in optical power. Setting the first terms on the right-hand sides of Eqs. 9 and 10 equal to each other and solving for the interocular difference in optical density yields the optical density difference that will achieve the anti-Pulfrich correction. Specifically,

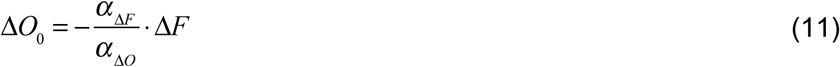

Negative values indicate that the transmittance of the left lens should be reduced to achieve an anti-Pulfrich correction. Positive values indicate that the transmittance of the right lens should be reduced.

### Summarizing effect sizes

To compare effect sizes across experiments and human observers, we report interocular delays in a particular condition estimated from the best-fit lines in each experiment, rather than using the raw PSE data itself. We used this approach for two reasons. First, this approach has the advantage of minimizing the effect of measurement error. Second, in experiments with blur differences (Exps. 1, 2, & 4), not all human observers collected data in identical conditions. Some human observers collected data with a maximum interocular difference in focus error (i.e. optical power) of ±1.5D while others collected data with a maximum difference of ±1.0D; some observers had difficulties performing the task with the larger difference. To compare interocular delays across human observers at the power difference associated with the most commonly prescribed monovision correction strength (i.e. 1.5D), we extrapolated using the best-fit lines to the data (Eq. 9).

## Results

Interocular differences in processing speed were measured in five separate experiments in each of four human observers. The same experimental paradigm was used to collect data across all five experiments. First, we describe the details of the procedure that was common across the experiments. Then, we describe each individual experiment. As a group, the experiments seek to establish: i) that contact lenses of different powers can induce the interocular mismatches in processing speed that underlie the reverse Pulfrich effect, ii) that anti-Pulfrich corrections with contact lenses are effective in eliminating the reverse Pulfrich, and iii) that interocular differences in image magnification have no impact on interocular differences in processing speed.

To measure interocular differences in processing speed, human observers collected data in a one-interval two-alternative forced-choice (2AFC) experiment. On each trial, observers viewed a dichoptically presented vertical target bar that oscillated horizontally in the frontal plane (Fig. 3A) while fixating a central dot (not shown). When the onscreen interocular delay is zero, onscreen disparity specifies that the target is moving in the plane of the screen. When the onscreen interocular delay is negative, the left-eye image onscreen trails the right-eye image onscreen, and onscreen disparity specifies that the target is following an elliptical trajectory outside the plane of the screen that is clockwise when viewed from above (‘front left’ motion). When the onscreen interocular delay is positive (i.e. an onscreen advance), the left-eye image onscreen leads the right-eye image onscreen, and onscreen disparity specifies that the target is following an elliptical trajectory outside the plane of the screen that is counter-clockwise when viewed from above (‘front right’ motion; Fig. 3B).

The task was to choose, on each trial, whether the target appeared to be undergoing ‘front right’ or ‘front left’ motion. In each condition, the proportion of times each observer chose ‘front right’ was plotted as a function of onscreen delay. This raw data was fit with a cumulative Gaussian function in each condition. Data and fits for the first human observer in the first experiment are shown in Fig. 3C. The point of subjective equality (PSE) indicates the onscreen delay necessary for the observer to report ‘front right’ on half of the trials (black arrows). This onscreen delay (or advance) of the left-eye onscreen image relative to the right-eye onscreen image is equal in magnitude and opposite in sign to the neural advance (or delay) induced by the image perturbations. Each experiment examines whether a particular interocular difference in image properties (i.e. a particular perturbation of the image in one eye) causes interocular differences in processing speed.

### Experiment 1: Contact Lenses

Experiment 1 measures the interocular differences in processing speed caused by blur differences induced by soft contact lenses having different powers. As mentioned earlier, contact lenses of different powers cause negligible interocular differences in magnification. Contact lenses thus isolate differences in optical blur from the possible confounding magnification differences caused by trial lenses (Fig. 4A). This experiment will determine whether the reverse Pulfrich effect occurs when interocular blur differences are not accompanied by interocular differences in magnification. It will also determine whether the reverse Pulfrich effect is caused by the most commonly used delivery system for monovision prescriptions.

**Figure 4.**
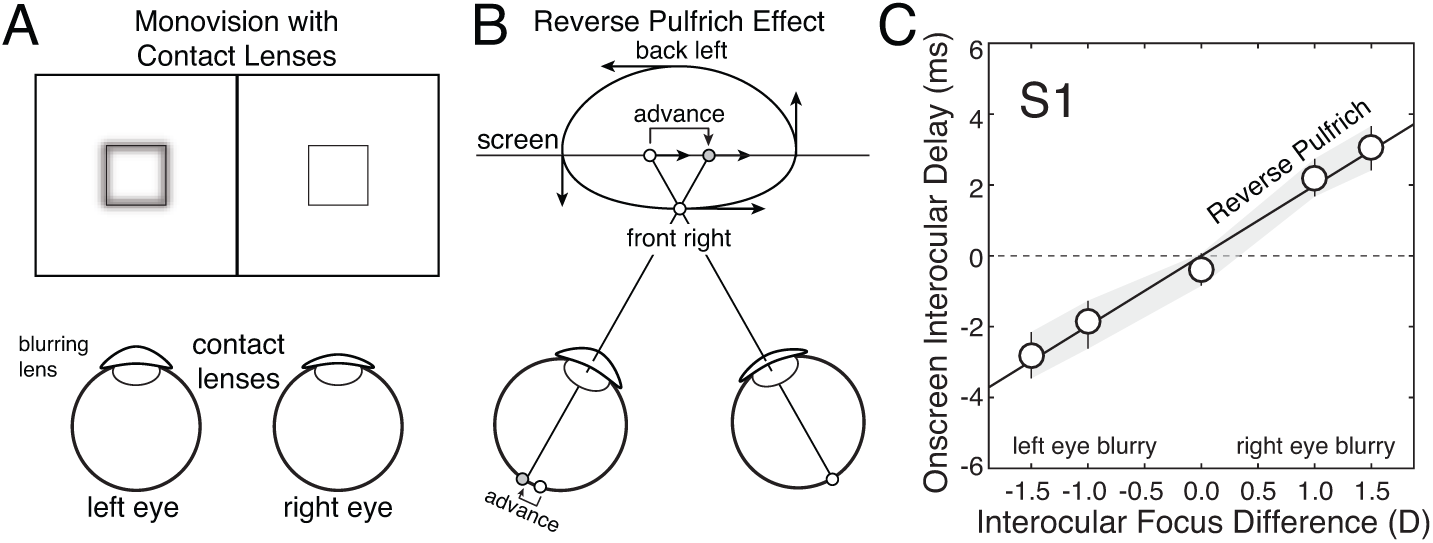
Reverse Pulfrich effect with contact lenses (Exp. 1). **A** Stimulus conditions with contact lenses. Contact lenses of different powers cause interocular differences in blur, but no differences in magnification. The differences in optical power (i.e. focus error) ranged from −1.5D to 1.5D, which are common monovision correction strengths. **B** The interocular difference in blur causes a mismatch in processing speed between the eyes—the blurrier image is processed more quickly—that leads to the reverse Pulfrich effect. Horizontal oscillating motion in the frontal plane is perceived as ‘front right’ elliptical motion in depth (i.e. counter-clockwise when viewed from above). **C** Onscreen interocular delays that are required to null neural differences in processing speed induced by differences in optical power between the eyes. Error bars indicate 68% confidence intervals from 1000 bootstrapped datasets.

One eye—the perturbed eye—was fit with a contact lens that blurred the stimulus (see Methods). The other eye was fit with a contact lens that sharply focused the stimulus. As expected, we found that contact lenses of different powers cause a reverse Pulfrich effect. When the left eye is blurred, a target stimulus oscillating in the frontal plane with no onscreen delay is incorrectly perceived as undergoing ‘front right’ motion in depth. The reverse Pulfrich effect occurs because the image in the blurrier eye is processed more quickly than the image in the sharper eye (Fig. 4B). To null this effect, the left eye must be delayed onscreen by an amount equal in magnitude but opposite in sign to the advance in neural processing speed.

Data from the first human observer is shown in Fig. 4C. When the left eye was at its blurriest and the right eye was sharp (i.e. Δ*F*=-1.5D), the left-eye stimulus had to be delayed onscreen by 2.8ms from baseline. When the left eye was sharp and the right eye was at its blurriest (i.e. Δ*F*=1.5D), the left-eye stimulus had to be advanced onscreen by 3.1ms from baseline. A similar pattern of results was found for all four human observers. Across observers, the blurrier eye was processed 1.9ms faster on average (SD=1.0ms).

These mismatches in processing speed imply that monovision corrections, which are most often delivered by contact lenses, can cause substantial misperceptions of motion (Burge et al., 2019). It may be advisable for optometrists and ophthalmologists to make their patients aware of these motion illusions when prescribing monovision, just as it is commonplace to motion the associated decrease in stereoacuity (Erickson & Schor, 1990; McGill & Erickson, 1988; Westheimer & McKee, 1980).

### Experiment 2: Trial Lenses

Experiment 2 measures the interocular differences in processing speed induced by trial lenses having different powers. As mentioned earlier, trial lenses with different powers cause non-negligible magnification differences—1.8% for a 1.5D difference—in addition to blur differences (Fig. 5A). Exp. 1 demonstrated that magnification differences are not necessary for the reverse Pulfrich effect to occur. But magnification differences could, in principle, interact with blur differences to strengthen or weaken the reverse Pulfrich effect. To examine this issue, we re-ran each human observer in conditions that were identical to those in Exp. 1, except that trial lenses, instead of contact lenses, induced the differences in optical blur.

**Figure 5.**
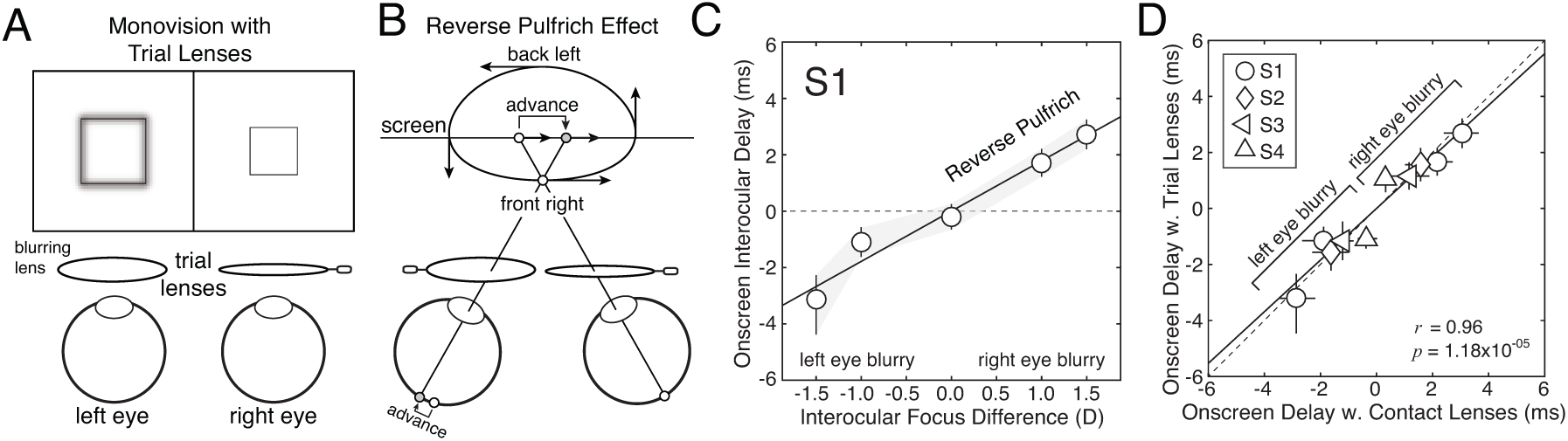
Reverse Pulfrich effect with trial lenses (Exp. 2). **A** Stimulus conditions with trial lenses. Trial lenses of different powers cause interocular differences in blur and in magnification. The differences in optical power (i.e. focus error) ranged from −1.5D to 1.5D. **B** The interocular difference in blur causes a mismatch in processing speed between the eyes—the blurrier image is processed more quickly—that leads to the reverse Pulfrich effect. Horizontal oscillating motion in the frontal plane is perceived as ‘front right’ elliptical motion in depth (i.e. counter-clockwise motion when viewed from above). **C** Onscreen interocular delays required to null neural delays induced by differences in optical power between the eyes. Error bars indicate 68% confidence intervals from 1000 bootstrapped datasets. **D** Onscreen interocular delays from trial lenses vs. contact lenses for each individual observer (symbols) in all conditions measured. Processing delays induced by contacts and trial lenses with equivalent power differences are nearly identical; the best-fit regression line has a slope of 0.92 (solid line). The interocular difference in magnification caused by the trial lenses has no effect on processing speed.

One eye—the perturbed eye—was fit with a trial lens that blurred the stimulus (see Methods). The other eye was fit with a trial lens that sharply focused the stimulus. The blurrier eye is processed more quickly, causing a reverse Pulfrich effect (Fig. 5B).

Data from the first human observer is shown in Fig. 5C. The onscreen interocular delay that is required to null the reverse Pulfrich effect changes linearly with the interocular difference in focus error, just as with contact lenses. In this observer, in the condition when the left eye was at its most blurry (Δ*F*=-1.5D), the left-eye image had to be delayed onscreen by 3.1ms. When the right eye was most blurry (Δ*F*=+1.5D), the left eye had to be advanced onscreen by 2.7ms. Again, a similar pattern of results was found for all human observers. Across observers, the blurrier eye was processed 2.1ms faster on average (SD=0.5ms). These findings replicate the primary result of Burge et al. (2019).

To check whether magnification differences had any influence on the size of the reverse Pulfrich effect, we plotted the effect size measured with trial lenses in each condition against the effect size measured with contact lenses in the same condition, for all observers (Fig. 5D). Trial lenses and contact lenses yielded very similar effect sizes that were tightly correlated across all human observers (*r*=0.96; *p*=1.18×10^−5^). The magnification differences caused by trial lenses, therefore, do not influence the size of the reverse Pulfrich effect caused by interocular blur differences.

### Experiment 3: Luminance Differences

Experiment 3 measures the interocular differences in processing speed caused by luminance differences between the eyes. This experiment is useful for two reasons. First, measuring the decrease in processing speed caused by darkening the image in one eye is necessary to test whether anti-Pulfrich monovision corrections are effective with contact lenses. Second, replicating results from the classic literature increases confidence that the current paradigm is producing valid results.

The image to one eye—the perturbed eye—was darkened onscreen by a factor equivalent to the transmittance of a neutral density filter with a particular optical density (Fig. 6A; see Methods). The other eye was left unperturbed. Both eyes were sharply focused on the stimulus. As expected, we found that a luminance difference between the eyes causes the classic Pulfrich effect; the darker image is processed more slowly. For a target stimulus oscillating in the frontal plane with no onscreen interocular delay, the percept is now of ‘front left’ motion in depth (Fig. 6B).

**Figure 6.**
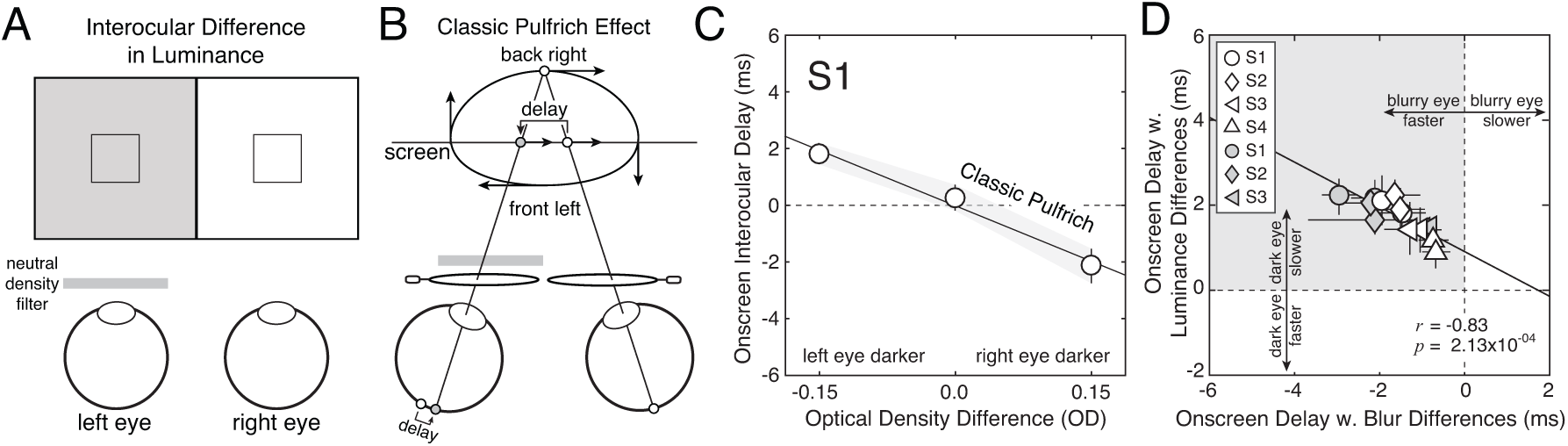
Classic Pulfrich effect with luminance differences (Exp. 3). **A** Stimulus conditions with interocular luminance differences. The image in one eye was darkened onscreen by a factor equal to the transmittance of a neutral density filter with a particular optical density; the other eye was left unperturbed. The differences in optical density ranged from −0.15OD to 0.15OD, corresponding to a 30% transmittance difference between the left and right eyes. **B** The luminance differences cause a mismatch in processing speed between the eyes—the darker image is processed more slowly. The classic Pulfrich effect results. Horizontal oscillating motion in the frontal plane is misperceived as ‘front left’ elliptical motion in depth (i.e. clockwise motion when viewed from above). **C** Onscreen interocular delays required to null the neural delays induced by interocular luminance differences. Error bars indicate 68% confidence intervals from 1000 bootstrapped datasets. **D** Interocular delays induced by luminance differences (i.e. |Δ*O*|=0.15OD) are plotted against interocular delays induced by blur differences (i.e. |Δ*F*|=1.0D) in individual observers from the current study (white symbols) and from Burge et al. (2019) (gray symbols). To isolate the factor of interest—the similarity of effect size due to interocular differences blur and luminance—we plot onscreen delays with respect to the perturbed eye rather than with respect to the left eye. In individual observers, the size of the reverse and classic Pulfrich effects are correlated (*r*=-0.83; *p*=2.13×10^−4^).

Data from the first human observer is shown in Fig. 6C. Just as with the reverse Pulfrich effect, the onscreen interocular delay required to null the classic Pulfrich effect changes linearly with the interocular difference in optical density. But the sign of the slope relating the difference is now negative instead of positive. When the left eye was darkest (i.e. Δ*O*=-0.15OD), the left-eye image had to be advanced onscreen by 1.8ms to null the neural delay. When the right eye was darkest (i.e. Δ*O*=+0.15OD), the left eye had to be delayed onscreen by 2.1ms to null the neural delay. Again, similar results were found for all observers; the darker eye was processed 1.6ms more slowly on average (SD=0.5ms).

Currently, it is unknown whether the interocular mismatches in processing speed induced by luminance and blur differences are mediated by common or partially shared neural mechanisms. To help constrain the answer to this question, we examine whether the sizes of the classic Pulfrich effect and the reverse Pulfrich effect were correlated amongst observers. (Note: to quantify the effect of blur differences on neural delay we averaged the reverse Pulfrich effect sizes for each observer across Exps. 1 and 2.) Fig. 6D plots the onscreen interocular advance (or delay) required to null the neural delay (or advance) caused by interocular differences in luminance and in blur.

Observers with large reverse Pulfrich effects tended to have large classic Pulfrich effects (*r* =-0.94, *p*=4.52×10^−4^). However, one must be careful not to place too much interpretative weight on a correlation computed from a very small number of observers. Accordingly, we included data from three additional observers from a previously published paper to increase the power of the analysis. Data from the current experiments are shown as white symbols. Data from Burge et al. (2019) is shown as gray symbols. Clearly, a similar trend is present in the previous dataset (*r* =-0.88; *p*=2.17×10^−2^). Across both datasets, the correlation was strong (*r* =-0.83; *p*=2.13×10^−4^). More data must be collected before drawing a firm conclusion, but the preliminary evidence suggests that the size of the reverse Pulfrich effect is correlated with the size of the classic Pulfrich effect in individual observers. If this preliminary evidence holds up, the result may be useful in attempts to understand the neurophysiological mechanisms that underlie these effects.

### Experiment 4: Anti-Pulfrich Corrections with Contact Lenses

Experiment 4 measures whether anti-Pulfrich monovision corrections delivered with contact lenses can eliminate the interocular differences in processing speed that cause the reverse Pulfrich effect. The logic of an anti-Pulfrich correction is simple. Decreasing luminance and increasing blur have opposite effects on processing speed; by tinting the blurring lens, it should be possible to simultaneously null the two effects for a large range of target distances (Fig. 7A; see Discussion). The efficacy of anti-Pulfrich monovision corrections has been demonstrated previously with trial lenses (Burge et al., 2019). Here, we show that anti-Pulfrich corrections work with contact lenses.

**Figure 7.**
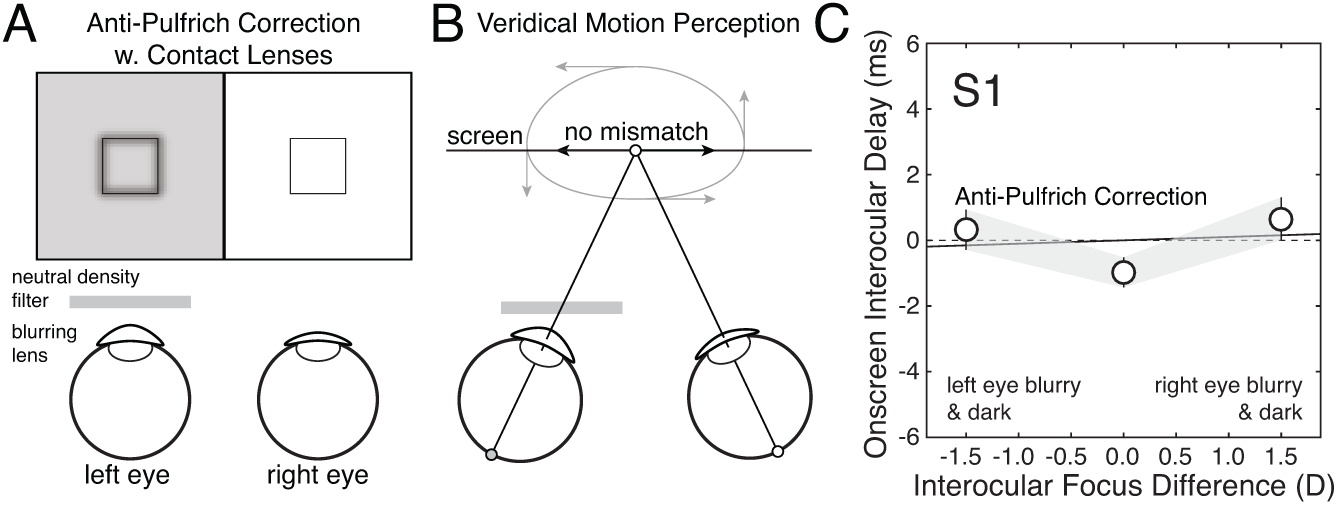
Anti-Pulfrich corrections with contact lenses null the reverse Pulfrich correction (Exp. 4). **A** Stimulus conditions for anti-Pulfrich monovision corrections. Darkening the image in the blurrier eye can eliminate the interocular differences in processing speed otherwise caused by blur. **B** Restoring the parity of processing speed eliminates the misperceptions associated with the reverse Pulfrich effect (dashed ellipse and arrows) and restores the veridical perception of moving objects (solid arrows). **C** Onscreen interocular delays are no longer required to null misperceptions of motion in depth, because anti-Pulfrich corrections (i.e. appropriately tinting the blurring lens) eliminates interocular differences in processing speed caused by blur alone. Error bars indicate 68% confidence intervals from 1000 bootstrapped datasets. Appropriately tinting the near contact lens in a pair of contact lenses delivering a monovision correction could eliminate the misperceptions of distance and 3D direction for far moving objects.

To determine the optical density of the filter that is appropriate to pair with a particular focus error, we compared how blur and luminance differences impacted processing speed in each human observer. The ratio of the slopes of the best-fit regression lines in Experiments 1 and 3 (see Figs. 4C, 6C, and Eq. 11) specifies the optical density required to null the change in processing speed due to a given blur difference. We found that appropriately darkening the blurry image successfully eliminates the mismatches in processing speed and restored veridical depth and motion perception (Fig. 7B).

Data from the first human observer is shown in Fig. 7C. The anti-Pulfrich correction with contact lenses was clearly successful at nulling the reverse Pulfrich effect in this observer. With an anti-Pulfrich correction, the interocular difference in processing speed that was caused in this observer by contact lenses with a 1.5D difference in optical power was reduced from 2.9ms to 0.1ms in this observer. Similarly successful results were obtained for all human observers. The average interocular difference in processing speed for a 1.5D difference in optical power was reduced from 1.9ms to −0.1ms (SD=0.3ms).

The first observer required the largest difference in optical density to null the reverse Pulfrich effect. Nulling the reverse Pulfrich effect for a 1.5D interocular difference in optical power required an interocular difference in optical density of 0.23OD. An optical density of 0.23OD corresponds to a transmittance of 59% (Eq. 4). Across observers, the required transmittance in the dark lens ranged from 59% to 89%. For reference, a standard pair of sunglasses transmits only 25% of the incoming light (i.e. optical density of 0.60). Thus, the required difference in transmittance required between the eyes for a successful anti-Pulfrich correction is rather slight (see Discussion).

### Experiment 5: Magnification Differences

Experiment 5 measures whether magnification differences between the images in the two eyes can cause processing speed differences. The results of Experiments 1 and 2 showed that magnification differences do not impact processing speed when differences in optical blur are present. The current experiment tests directly whether magnification differences can cause differences in processing speed when blur differences are absent.

Onscreen interocular differences in magnification were introduced, while both eyes were kept sharply focused and equally bright (Fig. 8A). The onscreen magnification difference (3.6%) was double the magnification difference caused by trial lenses that differ in power by 1.5D. Under these conditions, processing speed was equal in both eyes and motion perception was veridical; targets specified by disparity to be oscillating in the frontal plane were correctly perceived as oscillating in the frontal plane (Fig. 8B).

**Figure 8.**
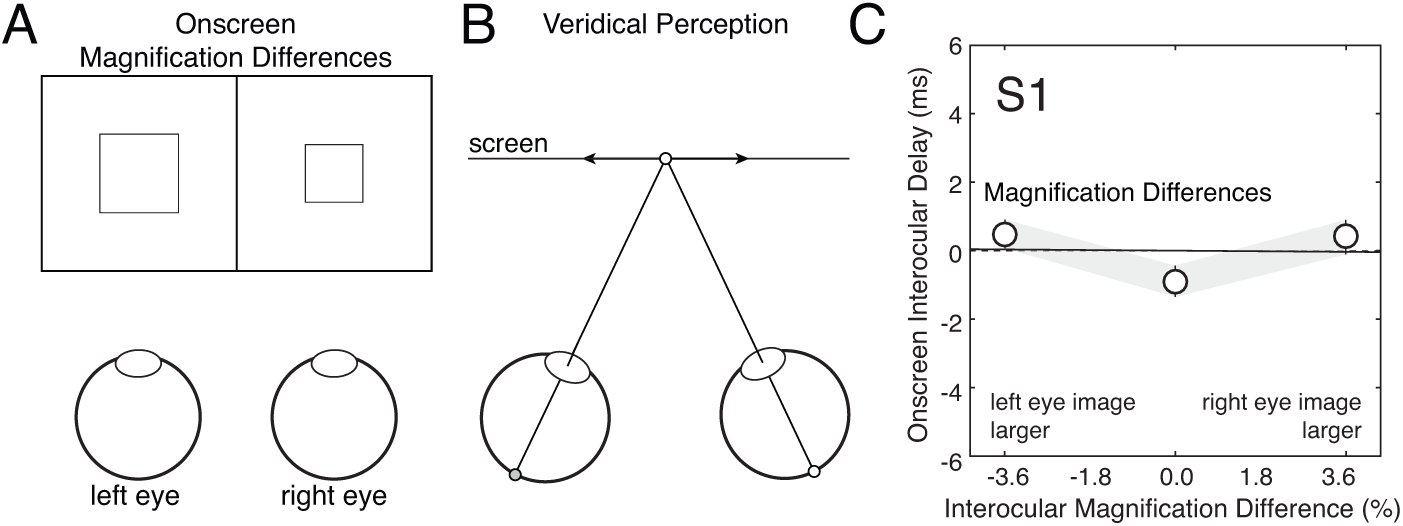
Magnification differences do not cause motion-in-depth misperceptions (Exp. 5). **A** Stimulus conditions with interocular differences in magnification. The image in one eye was larger than the image in the other eye. Both images were equally sharp and equally bright. **B** Magnification differences do not cause motion-in-depth misperceptions. Horizontally oscillating motion in the frontal plane is perceived veridically. **C** Onscreen interocular delays equal zero for all interocular differences in magnification. Error bars indicate 68% confidence intervals from 1000 bootstrapped datasets.

Data from the first human observer is shown in Fig. 8C. The largest magnification differences caused negligible interocular mismatches in processing speed. Similar results were obtained for all human observers. Across observers, the average interocular delay for a magnification difference of 3.6% was −0.0ms (SD=0.1ms). Magnification differences therefore do not cause the images in the two eyes to be processed at different speeds.

### Summary of Experimental Results

The pattern of results across experiments is remarkably consistent for all human observers (Fig. 9). In each observer, blur differences induced by contact lenses (Exp. 1) and trial lenses (Exp. 2) both cause the reverse Pulfrich effect; the image in the blurrier (i.e. perturbed) eye is processed faster than the image in the sharper eye. In each observer, luminance differences cause the classic Pulfrich effect (Exp. 3); the darker image is processed slower than the brighter image. In each human observer, anti-Pulfrich monovision corrections that are delivered with contact lenses eliminate the mismatches in processing speed that underlie the reverse Pulfrich effect (Exp. 4). And in each human observer, magnification differences cause no differences in processing speed between the eyes (Exp. 5). The consistency of these results across the four human observers in this report, and the similarity of these results to previously published findings, should increase confidence that these findings are solid and will be replicable in new populations.

**Figure 9.**
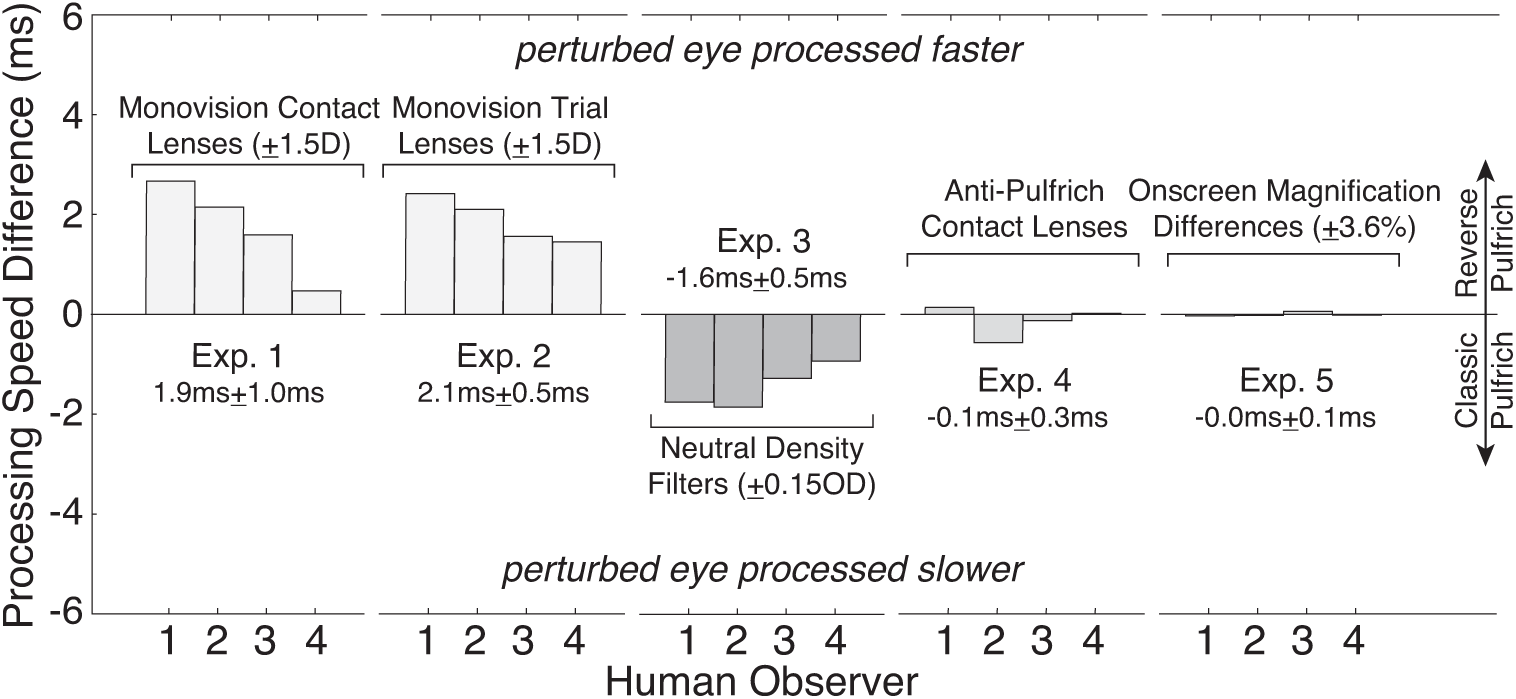
Summary of experimental data. Effect sizes across all experiments and human observers. Blurring one eye with contact lenses (Exp. 1) or blurring one eye with trial lenses (Exp. 2) causes the image in that same eye to be processed more quickly, leading to a reverse Pulfrich effect. Darkening one eye causes the image in that eye to be processed more slowly (Exp. 3), leading to a classic Pulfrich effect. Anti-Pulfrich corrections eliminate the increase in processing speed caused by blur alone by appropriately darkening the image in the blurry eye (Exp. 4), thereby eliminating the reverse Pulfrich effect. Interocular differences in magnification (Exp. 5) do not impact interocular differences in processing speed.

## Discussion

The reverse Pulfrich effect can be caused by blur differences that are induced by soft contact lenses. For 1.5D differences in optical blur, a common monovision correction strength, the blurrier image is processed faster by approximately 2ms. Under certain conditions, these small differences in processing speed may provoke large misperceptions of depth (see Fig. 1). Fortunately, anti-Pulfrich monovision corrections, which leverage the fact that increased blur and reduced retinal illuminance have opposite effects on processing speed, can eliminate the misperceptions in a large subset of viewing conditions (see below). These findings with soft contact lenses are quite likely to generalize to other approaches for delivering monovision corrections: semi-rigid contact lenses, surgically-implanted intraocular lenses, or refractive surgery. We have also demonstrated that magnification differences of similar magnitude to those induced by trial lenses (±1.5D) do not cause or modify the reverse Pulfrich effect. Together, these results place current explanations for the reverse Pulfrich effect on firm empirical grounds, invite a re-examination of monovision prescribing practices, and suggest potential directions for improving corrections for presbyopia.

### Measuring the reverse Pulfrich effect in the clinic

Many millions of people currently wear monovision corrections to compensate for presbyopia (Burge et al., 2019; Cope et al., 2015; Ingenito, 2015; Morgan et al., 2019). The prevalence of monovision corrections, the potential ramifications of the reverse Pulfrich effect (see Fig. 1), and the potential compensatory function of anti-Pulfrich corrections suggest a need for tests that can be deployed in the clinic.

There are two primary obstacles to developing tests for use in the clinic. The first obstacle is the development of cheap portable displays that can render stereo-3D content of sufficiently high spatial and temporal resolution so that useful measurements can be made. Ongoing work is attempting to address this issue (Vancleef et al., 2019). The second obstacle is that the time available to gather data in the clinic (e.g. minutes) is severely limited compared to the time available to gather data in the lab (e.g. hours). Thus, it is of paramount importance to develop methods that enable the rapid collection of high-quality data, that require little or no training, and that can be used with non-traditional populations including children. We are working to adapt target-tracking methods for continuous psychophysics for this purpose (Bonnen, Burge, Yates, Pillow, & Cormack, 2015).

### Anti-Pulfrich monovision corrections: potential and limitations

The results with anti-Pulfrich corrections suggest they have potential for clinical practice, but it is important to discuss their limitations and highlight the most important directions for future work. An effective anti-Pulfrich correction requires that a tint be applied to the lens forming the blurrier image, but the lens forming the blurrier image varies with target distance. For example, appropriately tinting the near lens will eliminate the reverse Pulfrich effect for far targets, but aggravate it for near targets. Thus, anti-Pulfrich corrections can only work for a subset of target distances. Assuming that tinting the near lens is the preferred solution—which is plausible because accurate perception of moving targets is probably more important for tasks at far than at near distances (e.g. driving vs. reading)—the range of distances for which motion misperceptions may be reduced or eliminated can be considerable: from the near point of the far lens to infinity. This range may be even larger for early presbyopes, who have some residual ability to accommodate, because they tend preferentially focus the far lens (Almutairi, Altoaimi, & Bradley, 2018). However, these issues clearly need further study.

Understanding the effect of ambient illumination changes on both the reverse and classic Pulfrich effects is critical to determining how practical anti-Pulfrich corrections would be in a real-world setting. The classic Pulfrich effect is known to increase with decreases in ambient illumination (Lit, 1949; Rogers & Anstis, 1972; Wilson & Anstis, 1969). It is unknown how light level affects the reverse Pulfrich effect. If decreases in ambient illumination change the sizes of the reverse and classic Pulfrich effects by the same amounts, anti-Pulfrich monovision corrections will be straightforward to implement. However, if the two effects change differently with light level, prescribing anti-Pulfrich corrections may be more challenging. If so, a given difference in optical blur would have to be compensated for by transmittance differences that change with light level. Fortunately, photochromic contact lens technologies present a possible solution (Molock, Cullerton, Spaulding, & Shivkumar, 2005).

Photochromic lenses reduce their transmittance with increases in ambient light levels. A photochromic anti-Pulfrich correction could be applied with a photochromic contact lens in one eye and a standard contact lens in the other eye. The photochromic properties of the contact lens could be tuned to compensate for the manner in which the effect sizes change with light level. It may also be possible to develop an anti-Pulfrich photochromic correction that operates when the user is outdoors, and that reverts to classic monovision when the user is indoors, where accurate perception of motion is likely to be less critical. Indoors, all the available light energy would be transmitted to the eye that is focused at near, which would benefit near visual tasks like reading. Another, perhaps simpler, approach might be to combine a classic monovision correction with sunglasses with custom transmittances in each eye that could provide an anti-Pulfrich correction when the patient is outdoors.

Finally, it would be important to determine whether the transmittance (e.g. tint) differences required for an anti-Pulfrich correction pose a cosmetic issue. In general, these differences in transmittance are small. For the observers of the current study, the required transmittance in the dark lens ranged from 59% to 89% of incoming light, assuming a 1.5D difference in optical power. A common pair of sunglasses transmits only 25% of the incoming light. It remains to be seen whether these transmittance differences would cause a cosmetic impediment for everyday wear. If so, we note that the issue is likely to occur only for anti-Pulfrich corrections prescribed with contact lenses, and not with surgically-implanted intraocular lenses (Davidson et al., 2016; Xiao et al., 2011). Contact lenses cover the iris and are visible, and intraocular lenses are inserted into the capsular bag and are generally not visible. However, before ophthalmologists consider surgically implanting anti-Pulfrich monovision intraocular lenses, significant further study is required.

### The impact of magnification differences on binocular processing

We have found no evidence that interocular differences in magnification cause interocular differences in processing speed. Contact lenses and trial lenses with equivalent power differences cause reverse Pulfrich effects of nearly identical size (Exp. 1 and Exp. 2; see Fig. 5D). Additionally, onscreen magnification differences that were unaccompanied by blur had no measurable impact on processing speed differences between the eyes. Magnification differences, however, do impact other aspects of binocular visual processing. As magnification differences increase, binocular contrast sensitivity worsens (Jiménez, Ponce, & Anera, 2004), binocular summation breaks down (Jiménez et al., 2004; Katsumi, Tanino, Hirose, 1986, 1986), fusion times increase, the largest disparity eliciting a depth percept decreases (Jiménez, Ponce, del Barco, Díaz, & Pérez-Ocón, 2002), and stereopsis functions less well (Highman, 1977; Lovasik & Szymkiw, 1985; Vlaskamp, Filippini, & Banks, 2009). Horizontal and vertical magnification differences caused by unilateral magnifiers (i.e. induced aniseikonia) can cause misperceptions of surface orientation (Ogle, 1950). Horizontal magnification differences have also been reported to cause Pulfrich-like effects (Ames, 1946). But these effects can be attributed to changes in the relative spatial positions of the target projections due to the prismatic properties of the magnifier, rather than to an induced interocular difference in processing speed (Miles, 1953). In other words, instead of a time delay causing a neural disparity for moving objects, the magnifier causes an actual disparity in the retinal images. The results reported in this manuscript indicate strongly that magnification differences do not impact the relative speed of processing between the eyes. Blur differences, not magnification differences, drive the reverse Pulfrich effect.

### Applicability of current results to the presbyopic population

The human observers tested in the current experiments were between the ages of 25 and 30 and were thus all non-presbyopic. However, due to the experimental design, these observers unable to clear induced optical blur with accommodation. In this respect, the non-presbyopic observers were like presbyopes in the current experiments. Still, it is unknown whether the current results will generalize to the population of presbyopes. Presbyopes are more likely to be adapted to how blurry retinal images appear. The inability to accommodate increases the likelihood that blurry images are formed on the retinas; images that were perceived as in best focus were consistent with the optics of the observers’ own eyes (Artal et al., 2004; Sawides, de Gracia, Dorronsoro, Webster, & Marcos, 2011). Whether this increased exposure to blur decreases (or increases) presbyopes’ susceptibility to the reverse Pulfrich effect is unknown. Future research will have to evaluate the prevalence and range of effect sizes in the normal and presbyopic populations.

### Stability of the reverse Pulfrich effect over time

Do the processing speed differences associated with the reverse Pulfrich effect decrease with extended exposure to differences in optical blur? The processing speed differences underlying the classic Pulfrich effect are known to decrease over an extended period of time, if the image in one eye is consistently darker than the image in the other eye (Rogers & Anstis, 1972; Standing, Dodwell, & Lang, 1968; Wolpert, Miall, Cumming, & Boniface, 1993). With the reverse Pulfrich effect, however, the eye with the blurrier image depends on the distance of the target being viewed, so one eye is unlikely to be consistently blurrier than the other. It is known that presbyopes that habitually wear monovision corrections are more likely to be adapted to the appearance of optical blur differences between the eyes (Radhakrishnan, Dorronsoro, Sawides, Webster, & Marcos, 2015). The same is likely to be true of non-presbyopes with mild anisometropia. But it is unknown whether adaptation to visual appearance is accompanied by an adaptation that decreases the processing speed differences underlying the reverse Pulfrich effect. Future work will be required to determine whether the reverse Pulfrich effect diminishes with extended exposure to moderate differences in optical power between the eyes.

### The reverse Pulfrich effect in the real world

To date, the reverse Pulfrich effect has been measured only with simple laboratory stimuli. Ultimately, it will be important to understand how the effect manifests with real world (i.e. natural) stimuli. One important facet of this understanding will be the ability to predict the interocular differences in processing speed from the image properties in the two eyes. This will be a difficult problem to solve. But recent developments in the ability to estimate the cues most relevant to the effect from natural images (i.e. defocus blur, binocular disparity, and motion) provide reason for optimism (Burge & Geisler, 2011; 2012; 2014; 2015). Models that compute cue values directly from images (‘image-computable models’) have found recent success in predicting human performance (Chin & Burge, 2020; Kane, Bex, & Dakin, 2011; Morgenstern et al., 2020; Schütt & Wichmann, 2017; Sebastian, Abrams, & Geisler, 2017). However, to our knowledge, there exists no theoretical or empirical work that tightly links the properties of natural images to processing speed. Methods for learning the most useful stimulus features for particular tasks may be helpful to these efforts (Burge & Jaini, 2017; Geisler, Najemnik, & Ing, 2009; Jaini & Burge, 2017). Recent research with simple stimuli, which will be helpful to these goals, has shown that processing speed is directly impacted by the spatial and spatial frequency properties of images (Burge et al., 2019; Lages, Mamassian, & Graf, 2003; Min, Reynaud, & Hess, 2020). But a computational theory that relates the properties of natural images to processing speed is necessary for a full scientific understanding of Pulfrich-related phenomena. Such a theory may prove useful for understanding any vision system (biological or machine) that must combine complementary streams of information that are processed with different speeds.

## Conclusions

The reverse Pulfrich effect can be caused by contact lenses delivering monovision corrections, and can be eliminated with contacts delivering anti-Pulfrich monovision corrections. Although many questions must be resolved before the suitability of anti-Pulfrich corrections can be determined for clinical practice, optometrists and ophthalmologists should consider making their patients aware of the reverse Pulfrich effect when prescribing monovision.

## Acknowledgments

The project that gave rise to these results received the support of a fellowship from “la Caixa” Foundation (ID 100010434; LCF/BQ/DR19/11740032) to V.R.L., Spanish Ministry of Science, Innovation, and Universities grants ISCIII DTS2016/00127, FIS2017-84753-R and CSIC LINKA20122 to C.D., and startup funds from the University of Pennsylvania to J.B..

## Author Contributions

V.R.L. collected and analyzed data, V.R.L. and J.B. wrote the paper, V.R.L., C.D., and J.B. conceived the project and edited the paper.

## Declaration of Interests

United States and International patents on anti-Pulfrich monovision corrections have been filed by the University of Pennsylvania and the Institute of Optics (CSIC) with J.B., V.R.L., and C.D. as inventors.

